# Performance assessment of age-adapted SOFA, qSOFA, and PELOD-2, PCIS, P-MODS for Hand, Foot and Mouth Disease

**DOI:** 10.1101/606574

**Authors:** Zhenjun Yu, Ali Li, Tingting Huang, Zebao He, Huazhong Chen, Jiansheng Zhu

**Affiliations:** Department of Infectious Diseases, Taizhou Enze Medical Center (Group) Enze Hospital, Taizhou, Zhejiang 318000, China; Department of Infectious Diseases, Affiliated Taizhou Hospital of Wenzhou Medical University, Linhai, Zhejiang 317000, China

**Author notes:** Corresponding author; E-mail address (J. Zhu).

**Keywords:** Hand, foot and mouth disease, Sequential organ failure assessment score, Pediatric logistic organ dysfunction score-2

## Abstract

**Objectives:** Hand, foot and mouth disease (HFMD) is a common infectious disease in children caused by intestinal virus and an important cause of child death. Early identification of critical HFMD and timely intervention are the key to reduce mortality. However, there is no available unified critical HFMD screening standard. This study aimed to explore the predictive evaluation of HFMD with critical illness scoring systems.

**Methods:** A total of 31 patients with mild HFMD, 30 with severe HFMD, and 25 with critical HFMD were included. The platelet index in age-adapted sequential organ failure assessment score (SOFA) was re-assigned to constitute the SOFA for HFMD (H-SOFA). The results of age-adapted SOFA, quick SOFA (qSOFA), and pediatric logistic organ dysfunction score-2 (PELOD-2), pediatric multiple organ dysfunction score (P-MODS), pediatric critical illness score (PCIS), H-SOFA of the three groups were compared.

**Results:** Significant differences in the following parameters were found between severe group and critical group: enterovirus 71 positive, heart rate, respiration, vomiting, cold sweat, moist rales, disturbance in consciousness, platelet, and blood glucose (P<0.05), as well as all critical illness scoring data (P<0.001). age-adapted SOFA, qSOFA, and PELOD-2, P-MODS, H-SOFA were positively correlated with critical HFMD (odds ratio (OR): 3.213, 8.66, 2.64, 2.56, and 4.297 respectively; P<0.01), with area under the curve (AUC) values of 0.938, 0.823, 0.848, 0.910, and 0.956, respectively. PCIS was negatively correlated with critical HFMD (OR=0.76, P<0.001), with an AUC value of 0.865.

**Conclusion:** Increase in platelet count was related to the severity of HFMD. Age-adapted SOFA, qSOFA, and PELOD-2, P-MODS, PCIS, H-SOFA had high predictive value on critical HFMD, with H-SOFA being the highest.

## Introduction

Hand, foot and mouth disease (HFMD) is a common infectious disease in children caused by intestinal virus, mainly occurring in children aged below 5 years. It is usually a self-limiting disease with a 7-to-10-day disease course and a good prognosis [1]. However, some patients with critical HFMD are prone to central nervous system (CNS) damage, which are manifested as aseptic meningitis, acute flaccid paralysis, encephalomyelitis, etc. Involvement of the brainstem can lead to autonomic nervous system (ANS) dysfunction, pulmonary edema myocardial damage, and even death; survivors often experience significant neurological sequelae [2].

In some critical HFMD cases, early identification difficulties and lack of timely and effective treatment are important causes of death. Therefore, early identification are the key to reduce the mortality. However, there is no available unified critical HFMD screening standard. According to the World Health Organization (WHO) guidelines, severe HFMD manifests itself in three distinct phases: CNS damage, ANS dysfunction, and later cardiopulmonary failure. Abnormal laboratory examinations, such as leukocytosis and thrombocytosis (>400×10^3^/ul), hyperglycemia, and abnormal imaging examination can indicate the diagnosis of severe HFMD [3]. According to the 2018 edition of HFMD guidelines of the National Health Commission of China (NHCC), the following indicators suggest that patients may develop severe HFMD: sustained high fever, neurological impairment, dyspnea, circulatory dysfunction, leukocytosis, hyperglycemia, and hyperlactatemia [4]. These indicators are partly based on the clinical experience and subjective judgment of doctors, and cannot be accurately quantified, hence affecting the clinical application.

In recent years, critical illness scoring systems based on quantitative classification have become an important clinical tool for the prediction and evaluation of the prognosis of critically ill patients and serve as a guide to clinical practice [5]. The commonly used severe disease scoring systems in children include pediatric logistic organ dysfunction score-2 (PELOD-2) [6], pediatric critical illness score (PCIS) [7], and pediatric multiple organ dysfunction score (P-MODS) [8]; application of these scores can help assess the severity of the disease, more accurately assesses the risk of death, and facilitate timely intervention. Sequential organ failure assessment score (SOFA) and quick SOFA (qSOFA) are widely used in the diagnosis and treatment of critical diseases [9]. However, both of them were developed for adults and hence are not suitable for children. Schlapbach et al. [10] designed the age-adapted SOFA and qSOFA based on the PELOD-2, and confirmed that age-adapted SOFA was superior to PELOD-2 and systemic inflammatory response syndrome score in evaluating critically ill children. Therefore, this article aimed to further study the effect of these scoring systems on the prediction and evaluation of critical HFMD.

## Methods

### Study participants

From January 2012 to June 2018, 2,832 children with HFMD were admitted to Affiliated Taizhou Hospital of Wenzhou Medical University and Taizhou Enze Medical Center (Group) Enze Hospital. A total of 31 patients with mild HFMD, 30 with severe HFMD, and 25 with critical HFMD were randomly included, and their clinical data were recorded in detail. Cases of hospitalization or death that occurred less than 1 day after hospitalization were excluded. Based on the 2017 WHO guidelines for HFMD [3] and the 2018 Chinese guidelines for HFMD [4], the included patients were divided into three groups: mild group, severe group, and critical group.

Mild group: patients presented with fever and erythra on hands, feet, mouth, or hips, without damage to the CNS or cardiopulmonary system. Severe group: based on the symptoms of mild cases, patients presented with CNS damage, without ANS dysfunction or cardiopulmonary failure. Critical group: based on the symptoms of severe cases, patients presented with ANS dysfunction or cardiopulmonary failure.

### Symptoms and signs

The general conditions, symptoms, and signs of the three groups were compared and analyzed, including the vital signs, fever days, hospitalization days, cough, hyperarousal, limb jitter, vomit, cold sweat, moist rales, convulsions, level of consciousness, and distribution of erythra.

### Laboratory parameters

Thirty-seven clinical laboratory parameters were detected and analyzed, including blood tests performed using an AutoAnalyzer (Seemed-2100, Sysmex Corp., Japan), blood coagulation detected by hematology analyzers (STA Compact, Stago Corp., France), biochemical and immune examination performed using an AutoAnalyzer (Architect ci16200, Abbott Corp., USA), and enteroviral etiology detected using PCR analyzer (ABI7300, Applied Biosystems Inc., USA). The detailed laboratory parameters were white blood cell count (WBC), neutrophil count (NEUT), lymphocyte count (LYMP), monocyte count (MONO), eosinophil count (EO), basophil count (BASO), hemoglobin (HB), hematocrit (HCT), platelet (PLT), prothrombin time (PT), activated partial thromboplastin time (APTT), fibrinogen (Fib), immunoglobulin G (IgG), immunoglobulin A (IgA), immunoglobulin M (IgM), C-reactive protein (CRP), procalcitonin (PCT), aminotransferase (ALT), aspartate transaminase (AST), alkaline phosphatase (ALP), glutamyl transpeptidase (GGT), total bilirubin (TB), direct Bilirubin (DB), indirect Bilirubin (IB), albumin (ALB), prealbumin (PB), sodium (Na), serum potassium (K), serum chloride (Cl), serum calcium (Ca), uric acid (UA), urea, serum creatinine (SCr), blood glucose (GLU), creatine kinase (CK), MB isoenzyme of creatine kinase (CKMB), lactate dehydrogenase (LDH), Enterovirus A group 71 (EV-A71), and common enterovirus type.

### Age-adapted SOFA and qSOFA

Schlapbach et al. [10] revealed that the evaluation of circulatory system and renal function, which was previously conducted based on the SOFA scale, was recently performed using the PELOD-2 rating scale, which includes the assignment of mean arterial pressure (MAP) and SCr, as age-adapted SOFA. Meanwhile, the age-adapted qSOFA was designed according to the 2005 adjusted diagnostic criteria for pediatric sepsis, in which the age-specific respiratory rate and systolic pressure were used to determine the score.

### H-SOFA

According to our previous studies, the increase of PLT level is associated with severe HFMD, which is also consistent with the opinion of WHO guidelines [3]. The values of coagulation system in the SOFA were considered due to the decline in PLT level, which was not conducive to the evaluation of HFMD. Therefore, the PLT levels (×10^3^/ul) were scored as follows: 150–299, 0 points; 100–149 or 300–399, 1 point; and 50–99 or 400–599, 2 points, 20–49 or 600–899), 3 points; and <20 or ≥900, 4 points. According to the above scoring principles, the special SOFA for HFMD (H-SOFA) was constituted.

The detailed assignment instructions of age-adapted SOFA, qSOFA, and H-SOFA described above can be found in the supplementary materials.

### PELOD-2, PCIS, and P-MODS

The PELOD-2, PCIS, and P-MODS were specially designed for children. Detailed scoring instructions can be found in the supplementary materials. Six scoring systems were included in this study, and the HFMD cases of the three groups were scored and compared.

### Statistical analysis

Statistical analysis was performed using SPSS version 22.0 (IBM Inc., USA). The tables were created using Microsoft^®^ Excel^®^ 2016 (Microsoft Corp., USA). Continuous variables were presented as a mean ± standard deviation. One way analysis of variance was used for comparison between groups. For further pairwise comparison, LSD-t test was applied for parameters with equal variance, and Tamhane’s T2 test was applied for parameters with different variance. The frequencies and percentages were given for qualitative variables. Significant differences were evaluated using chi-square test, and Fisher’s exact test was used when the numbers were too small. Binary logistic regression analysis was used to identify the scoring systems that correlated with critical HFMD. The predicting values were tested with receiver operating characteristic (ROC) curves, and quantified by calculating the area under the curve (AUC) and the 95% confidence interval. A two-tailed P value of <0.05 was considered significant.

## Results

No statistical differences were observed among three groups in terms of age, height, weight, sex ratio, and maximum body temperature (P>0.05), while vital signs such as heart rate, respiration, and MAP were significantly different (P<0.01). The etiology tests showed that EV-A71 infection rate was 0 in the mild group, 30.0% in the severe group and 64.0% in the critical group, with significant differences among the three groups (P<0.001) (Table 1). The comparison of symptoms and signs among the three groups showed that the symptoms like cough and distribution of erythra had no significant differences, but other signs such as hyperarousal, limb jitter, vomit, cold sweat, moist rales, convulsions, and disturbance in consciousness were significantly different (P<0.01) (Table 2).

**Table 1:** Comparison of general conditions, clinical characteristics, and pathogenic characteristics among the three groups

| Characteristics | Mild group<br>(n=31) | Severe group<br>(n=30) | Critical group<br>(n=25) | F/$\chi^2$<br>value | P value |
| --- | --- | --- | --- | --- | --- |
| Age, months | 24.03±18.95 | 31.37±22.47 | 24.28±13.94 | 1.408 | 0.251 |
| Height, cm | 86.18±14.74 | 90.13±15.50 | 84.00±11.72 | 1.336 | 0.269 |
| Weight, kg | 13.14±4.83 | 13.59±4.52 | 12.24±3.32 | 0.676 | 0.511 |
| Male/Female, n | 19/12 | 21/9 | 17/8 | 0.564 | 0.754 |
| Fever, days | 2.55±1.18 | 3.57±1.74 | 5.44±3.11 | 13.462 | <0.001 |
| hospitalization, days | 4.23±1.18 | 5.80±1.35 | 8.82±7.16 | 9.354 | <0.001 |
| Temperature, °C | 39.16±0.80 | 39.17±0.75 | 39.45±0.51 | 1.451 | 0.240 |
| Heart rate, bpm | 121.52±11.05 | 148.20±17.19 | 174.12±23.56 | 62.524 | <0.001 |
| Respiration, bpm | 25.87±4.50 | 37.87±10.77 | 53.92±15.67 | 45.809 | <0.001 |
| MAP, mmHg | 83.10±11.46 | 71.73±4.95 | 72.37±12.96 | 7.799 | 0.001 |
| EV-A71 infection, n (%) | 0 (0%) | 9 (30.0%) | 16 (64.0%) | 27.511 | <0.001 |

Abbreviations: bpm, beats per minute; MAP, mean arterial pressure; EV-A71, enterovirus A group 71.

**Table 2:** Comparison of clinical symptoms and signs among the three groups

| Symptoms and signs | Mild group<br>(n=31) | Severe group<br>(n=30) | Critical group<br>(n=25) | $\chi^2$<br>value | P<br>value |
| --- | --- | --- | --- | --- | --- |
| Cough, n (%) | 8 (25.8%) | 11 (36.7%) | 8 (32.0%) | 0.841 | 0.657 |
| Hyperarousal, n (%) | 3 (9.7%) | 22 (73.3%) | 22 (88.0%) | 40.741 | <0.001 |
| Limb jitter, n (%) | 7 (22.6%) | 20 (66.7%) | 22 (88.0%) | 25.925 | <0.001 |
| Vomit, n (%) | 5 (16.1%) | 14 (46.7%) | 14 (56.0%) | 10.644 | 0.005 |
| Cold sweat, n (%) | 0 (0%) | 0 (0%) | 8 (32.0%) | 17.271 | <0.001 |
| Moist Rales, n (%) | 0 (0%) | 1 (3.3%) | 11 (44.0%) | 22.863 | <0.001 |
| Convulsion, n (%) | 0 (0%) | 5 (16.7%) | 6 (24.0%) | 8.864 | 0.008 |
| Disturbance in consciousness,<br>n (%) | 0 (0%) | 1 (3.3%) | 11 (44.0%) | 22.863 | <0.001 |
| Hand with erythra | 30 (96.8%) | 26 (86.7%) | 22 (88.0%) | 2.273 | 0.373 |
| Foot with erythra | 29 (93.5%) | 28 (93.3%) | 24 (96.0) | 0.393 | 0.991 |
| Mouth with erythra | 30 (96.8%) | 29 (96.7%) | 23 (92.0%) | 1.015 | 0.677 |
| Hip with erythra | 16 (51.6%) | 16 (53.3%) | 16 (64.0%) | 0.976 | 0.614 |

By comparing the laboratory parameters of the three groups, significant differences were found in the WBC, NEUT, PLT, GLU, PT and APTT (P<0.01). By contrast, other parameters, such as LYMP, MONO, EO, BASO, HB, HCT, IgG, IgA, IgM, CRP, PCT, ALT, AST, ALP, GGT, TB, DB, IB, ALB, PB, K, Na, Cl, Ca, SCr, Urea, UA, CK, CKMB, LDH, Fib, had no significant differences (P>0.5) (Table 3). Data of the scoring systems of the three groups were compared, and significant differences were found in age-adapted SOFA, qSOFA, and PELOD-2, PCIS, P-MODS, H-SOFA (P<0.001) (Table 4).

**Table 3:** Comparison of laboratory parameters among the three groups

| Laboratory parameters<br>mean ± SD | Mild group<br>(n=31) | Severe group<br>(n=30) | Critical group<br>(n=25) | F<br>value | P<br>value |
| --- | --- | --- | --- | --- | --- |
| WBC | 10.32±4.05 | 13.07±4.72 | 15.60±7.61 | 6.411 | 0.003 |
| NEUT | 5.55±3.40 | 9.05±4.93 | 9.96±6.51 | 6.137 | 0.003 |
| LYMP | 3.74±2.40 | 3.13±1.58 | 4.72±3.32 | 2.795 | 0.067 |
| MONO | 0.77±0.41 | 0.71±0.43 | 0.94±0.60 | 1.648 | 0.199 |
| EO | 0.04±0.04 | 0.02±0.05 | 0.02±0.05 | 1.153 | 0.321 |
| BASO | 0.03±0.02 | 0.01±0.01 | 0.02±0.02 | 2.887 | 0.061 |
| HB | 120.68±10.29 | 115.60±8.16 | 116.24±19.74 | 1.319 | 0.273 |
| HCT | 0.36±0.03 | 0.34±0.02 | 0.35±0.06 | 1.324 | 0.272 |
| PLT | 289.00±88.43 | 314.67±103.32 | 371.04±101.50 | 5.007 | 0.009 |
| IgG | 7.98±3.73 | 11.66±6.51 | 11.21±7.71 | 2.836 | 0.065 |
| IgA | 0.43±0.27 | 0.62±0.52 | 0.43±0.24 | 2.198 | 0.119 |
| IgM | 1.12±0.46 | 1.27±0.40 | 1.23±0.48 | 0.788 | 0.459 |
| CRP | 15.14±21.38 | 14.71±34.46 | 5.02±4.69 | 1.265 | 0.288 |
| PCT | 0.15±0.15 | 0.26±0.42 | 0.32±0.68 | 0.348 | 0.708 |
| ALT | 18.13±10.17 | 15.73±4.11 | 18.68±12.12 | 0.819 | 0.444 |
| AST | 36.74±8.11 | 38.28±15.12 | 46.13±27.70 | 2.048 | 0.136 |
| ALP | 202.81±49.15 | 227.17±61.64 | 222.71±58.79 | 1.452 | 0.240 |
| GGT | 12.13±3.39 | 10.41±2.78 | 16.19±17.88 | 2.332 | 0.104 |
| TB | 6.97±1.79 | 6.61±2.48 | 6.14±3.91 | 0.617 | 0.542 |
| DB | 2.39±1.02 | 2.73±1.18 | 2.20±1.30 | 1.363 | 0.262 |
| IB | 4.56±1.03 | 3.70±1.36 | 3.55±2.64 | 2.878 | 0.062 |
| ALB | 45.35±3.32 | 45.76±2.86 | 43.54±5.83 | 2.038 | 0.137 |
| PB | 13.40±3.28 | 15.37±3.12 | 14.99±3.37 | 2.991 | 0.056 |
| K | 4.43±0.38 | 4.51±0.56 | 4.23±0.53 | 2.324 | 0.113 |
| Na | 137.82±2.71 | 137.50±2.86 | 136.13±3.58 | 2.318 | 0.105 |
| Cl | 101.19±3.64 | 101.58±2.05 | 100.72±4.81 | 0.390 | 0.678 |
| Ca | 2.51±0.31 | 2.46±0.12 | 2.39±0.18 | 2.134 | 0.125 |
| SCr | 35.20±6.48 | 29.03±12.72 | 29.27±18.24 | 2.104 | 0.129 |
| Urea | 3.43±0.89 | 3.82±0.79 | 3.94±1.40 | 1.875 | 0.160 |
| UA | 261.36±86.58 | 231.24±74.57 | 275.71±99.79 | 1.786 | 0.175 |
| GLU | 5.02±1.13 | 5.94±1.64 | 7.26±2.58 | 10.501 | <0.001 |
| CK | 103.90±54.14 | 168.77±248.95 | 155.20±235.25 | 0.886 | 0.416 |
| CKMB | 26.58±10.95 | 34.89±18.39 | 32.54±16.04 | 1.903 | 0.156 |
| LDH | 345.80±130.68 | 314.20±140.09 | 288.96±107.86 | 1.364 | 0.261 |
| PT | 11.97±0.99 | 14.44±0.76 | 14.77±1.71 | 12.387 | <0.001 |
| APTT | 27.32±2.72 | 35.63±3.19 | 38.43±5.58 | 17.911 | <0.001 |
| Fib | 2.89±0.84 | 3.45±0.28 | 3.56±0.95 | 2.057 | 0.141 |

Abbreviations: WBC, white blood cell count; NEUT, neutrophils count; LYMP, lymphocytes count; MONO, monocytes count; EO, eosinophils count; BASO, basophils count; HB, hemoglobin; HCT, hematocrit; PLT, platelet; IgG, immunoglobulin G; IgA, immunoglobulin A; IgM, immunoglobulin
M; CRP, C-reactive protein; PCT, procalcitonin; ALT, aminotransferase; AST, aspartate transaminase; ALP, alkaline phosphatase; GGT, glutamyl transpeptidase; TB, total bilirubin; DB, direct bilirubin; IB, indirect bilirubin; ALB, albumin; PB, prealbumin; Na, sodium; K, serum potassium; Cl, serum chloride; Ca, serum calcium; UA, uric acid; SCr, serum creatinine; GLU, blood glucose; CK, creatine kinase; CKMB, MB isoenzyme of creatine kinase; LDH, lactate dehydrogenase; PT, prothrombin time; APTT, activated partial thromboplastin time; Fib, fibrinogen.

**Table 4:** Comparison of the scores of each scoring system among the three groups

| Scoring systems<br>mean ± SD | Mild group<br>(n=31) | Severe group<br>(n=30) | Critical group<br>(n=25) | F value | P<br>value |
| --- | --- | --- | --- | --- | --- |
| Age-adapted SOFA | 0.06±0.25 | 1.80±1.30 | 6.00±2.86 | 84.774 | <0.001 |
| Age-adapted qSOFA | 0.03±0.18 | 1.73±0.58 | 2.60±0.65 | 193.73 | <0.001 |
| PELOD-2 | 0.26±0.68 | 1.40±0.86 | 4.96±3.35 | 44.197 | <0.001 |
| PCIS | 89.61±1.20 | 83.00±5.16 | 74.00±6.40 | 77.782 | <0.001 |
| P-MODS | 1.19±.133 | 1.77±1.48 | 5.40±2.23 | 48.886 | <0.001 |
| H-SOFA | 0.61±0.67 | 2.47±1.33 | 7.28±2.99 | 97.871 | <0.001 |

Abbreviations: SOFA, Sequential Organ Failure Assessment Score; qSOFA, quick SOFA; PELOD-2, Pediatric Logistic Organ Dysfunction Score-2; PCIS, Pediatric Critical Illness Score; P-MODS, Pediatric Multiple Organ Drgan Dysfunction Score; H-SOFA, Age-adapted SOFA for Hand Foot and Mouth Disease.

The indicators with statistical differences among the three groups were further compared and analyzed between the severe and critical groups. Significant differences were found in the following parameters: EV-A71 infection, heart rate, respiration, cold sweat, moist rales, disturbance in consciousness, PLT, and GLU (P<0.05). By contrast, no significant difference was observed in other parameters, like MAP, hyperarousal, limb jitter, vomit, convulsion, WBC, NEUT, PT, and APTT (Table 5). In addition, there were significant differences (P<0.001) in data comparison of all scoring systems between severe group and critical group (Table 5).

**Table 5:** Comparison of parameters and scoring systems between theseveregroup and critical group

| Etiology/symptoms and signs | F/$\chi^2$ value | P value | Laboratory parameters/scoring systems | F value | P value |
| --- | --- | --- | --- | --- | --- |
| EV-A71 infection | 6.358 | 0.012 | WBC | 1.447 | 0.398 |
| Heart rate | 4.578 | <0.001 | NEUT | 0.573 | 0.920 |
| Respiration | 4.337 | <0.001 | PLT | 2.132 | 0.036 |
| MAP | 0.234 | 0.994 | GLU | 2.675 | 0.009 |
| Hyperarousal | 1.832 | 0.181 | PT | 0.523 | 0.603 |
| Limb jitter | 3.439 | 0.064 | APTT | 1.369 | 0.178 |
| Vomit | 0.475 | 0.491 | Age-adapted SOFA | 6.791 | <0.001 |
| Cold sweat | 14.278 | <0.001 | Age-adapted qSOFA | 5.178 | <0.001 |
| Moist rales | 13.221 | <0.001 | PELOD-2 | 5.178 | <0.001 |
| Convulsion | 0.458 | 0.498 | PCIS | 5.663 | <0.001 |
| Disturbance in consciousness | 13.221 | <0.001 | P-MODS | 6.956 | <0.001 |
|  |  |  | H-SOFA | 7.452 | <0.001 |

Abbreviations: EV-A71, enterovirus A group 71; MAP, mean arterial pressure; WBC, white blood cell count; NEUT, neutrophils count; PLT, platelet; GLU, blood glucose; PT, prothrombin time; APTT, activated partial thromboplastin time; SOFA, Sequential Organ Failure Assessment Score; qSOFA, quick SOFA; PELOD-2, Pediatric Logistic Organ Dysfunction Score-2; PCIS, Pediatric Critical Illness Score; P-MODS, Pediatric Multiple Organ Dysfunction Score; H-SOFA, Age-adapted SOFA for Hand Foot and Mouth Disease.

Further logistic regression analysis showed that age-adapted SOFA, qSOFA, PELOD-2, P-MODS, H-SOFA, and critical HFMD were positively correlated (odds ratio (OR): 3.213, 8.66, 2.64, 2.56, and 4.297, respectively; P<0.01), and PCIS was negatively correlated with critical HFMD (OR: 0.76; P<0.001) (Table 6). ROC curve analysis showed that the AUC of age-adapted SOFA, qSOFA, and PELOD-2, P-MODS, H-SOFA were 0.938, 0.823, 0.848, 0.910, and 0.956 respectively, and that of PCIS was 0.865 (Table 7, Figure 1). Compared with other scoring systems, the redesigned H-SOFA had the highest AUC value, with a sensitivity of 88.0% and a specificity of 90.0%. age-adapted SOFA had the highest sensitivity of 96.0%, but its specificity was only 80.0%. By contrast, age-adapted qSOFA had the highest specificity of 93.33%, but its sensitivity was only 68.0% (Table 7, Figure 1).

**Table 6:** Logistic regression analysis of correlations between thescoring systems and critical HFMD

| Scoring systems | B | SE | Wald | df | P value | OR | 95% CI for OR |  |
| --- | --- | --- | --- | --- | --- | --- | --- | --- |
|  |  |  |  |  |  |  | Lower | Upper |
| Age-adapted SOFA | 1.17 | 0.32 | 13.36 | 1 | <0.001 | 3.213 | 1.72 | 6.01 |
| Age-adapted qSOFA | 2.16 | 0.59 | 13.63 | 1 | <0.001 | 8.66 | 2.75 | 27.23 |
| PELOD-2 | 0.97 | 0.30 | 10.79 | 1 | 0.001 | 2.64 | 1.48 | 4.71 |
| PCIS | 0.27 | 0.07 | 14.11 | 1 | <0.001 | 0.76 | 0.66 | 0.88 |
| P-MODS | 0.94 | 0.24 | 15.74 | 1 | <0.001 | 2.56 | 1.61 | 4.07 |
| H-SOFA | 1.46 | 0.41 | 12.38 | 1 | <0.001 | 4.297 | 1.91 | 9.68 |

Abbreviations: B, Independent variable coefficient; SE, standard error; df, degrees of freedom; OR, odds ratio; CI, confidence interval; SOFA, Sequential Organ Failure Assessment Score; qSOFA, quick SOFA; PELOD-2, Pediatric Logistic Organ Dysfunction Score-2; PCIS, Pediatric Critical Illness Score; P-MODS, Pediatric Multiple Organ Dysfunction Score; H-SOFA, Age-adapted SOFA for Hand Foot and Mouth Disease.

**Table 7:** ROC curve was applied to analyze the predictive value of each scoring system for critical HFMD

| Scoring systems | AUC | P value | Cut-off value | Sensitivity | Specificity | 95% CI for AUC |  |
| --- | --- | --- | --- | --- | --- | --- | --- |
|  |  |  |  |  |  | Lower | Upper |
| Age-adapted SOFA | 0.938 | <0.001 | 2 | 96.00 | 80.00 | 0.839 | 0.985 |
| Age-adapted qSOFA | 0.823 | <0.001 | 2 | 68.00 | 93.33 | 0.696 | 0.912 |
| PELOD-2 | 0.848 | <0.001 | 1 | 80.00 | 80.00 | 0.726 | 0.931 |
| PCIS | 0.865 | <0.001 | 76 | 80.00 | 86.67 | 0.746 | 0.942 |
| P-MODS | 0.910 | <0.001 | 2 | 92.00 | 76.67 | 0.802 | 0.970 |
| H-SOFA | 0.956 | <0.001 | 4 | 88.00 | 90.00 | 0.864 | 0.993 |

Abbreviations: ROC, receiver operating characteristic; AUC, area under the ROC curve; CI, confidence interval; SOFA, Sequential Organ Failure Assessment Score; qSOFA, quick SOFA; PELOD-2, Pediatric Logistic Organ Dysfunction Score-2; PCIS, Pediatric Critical Illness Score; P-MODS, Pediatric Multiple Organ Drgan Dysfunction Score; H-SOFA, Age-adapted SOFA for Hand Foot and Mouth Disease.

**Figure 1:**
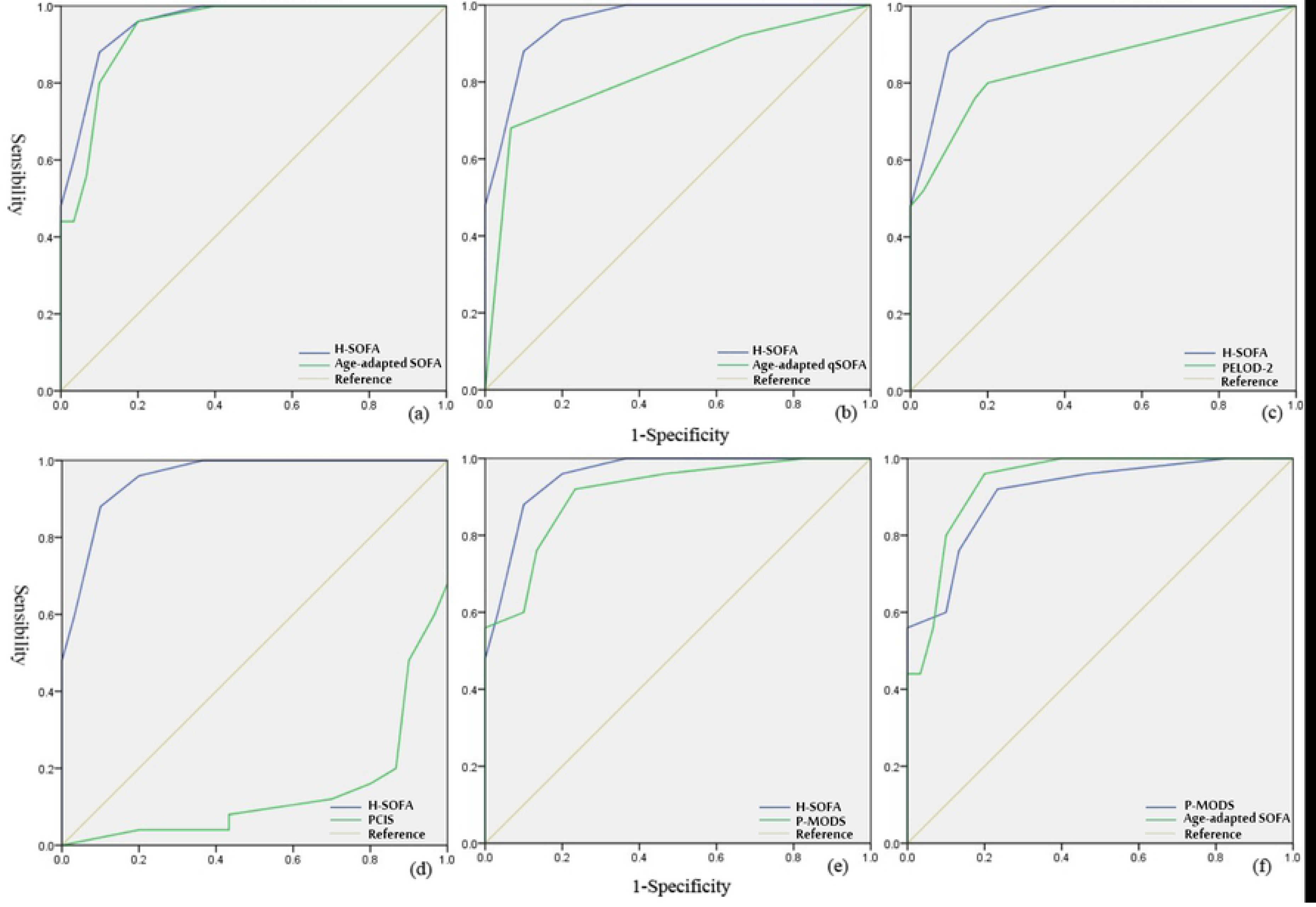
ROC curve comparisons among the scoring systems. a, the AUC of H-SOFA and age-adapted SOFA were 0.956 and 0.938; b, the AUC of H-SOFA and age-adapted qSOFA were 0.956 and 0.823; c, the AUC of H-SOFA and PELOD-2 were 0.956 and 0.848; d, the AUC of H-SOFA and PCIS were 0.956 and 0.865; e, the AUC of H-SOFA and P-MODS were 0.956 and 0.910; f, the AUC of P-MODS and age-adapted SOFA were 0.910 and 0.938.

Abbreviations: ROC, receiver operating characteristic; AUC, area under the ROC curve; SOFA, Sequential Organ Failure Assessment Score; qSOFA, quick SOFA; PELOD-2, Pediatric Logistic Organ Dysfunction Score-2; PCIS, Pediatric Critical Illness Score; P-MODS, Pediatric Multiple Organ Drgan Dysfunction Score; H-SOFA, Age-adapted SOFA for Hand Foot and Mouth Disease.

## Discussion

In this study, significant differences were observed among mild, severe, and critical patients with HFMD, mainly including heart rate, respiration, blood pressure, and symptoms or signs related to nervous system damage, such as hyperarousal, limb jitter, convulsion, and disturbance in consciousness. The severity of HFMD was not correlated with the symptoms like cough or distributions of erythra. This is also consistent with the definition of severe and critical HFMD proposed in the WHO guidelines [3,4]. However, the laboratory parameters that were significantly different among the three groups were only WBC, NEUT, PLT, GLU, PT, and APTT levels. Inflammatory indicators such as CRP and PCT, as well as liver or renal function-related indicators were not correlated with the severity of HFMD. HFMD is caused by infection of intestinal virus; severe or critical cases were associated with neurological damage, as well as neurogenic pulmonary edema, brainstem encephalitis, cardiopulmonary failure, etc. [11]. So liver and kidneys are not directly target organs, and severe liver or kidney damage can be seen in patients with critical HFMD who developed multiple organ dysfunction.

Consistent with previous studies, most of the children with CNS damage were infected with EV-A71 [12], and this study also showed that EV-A71 infection was the main cause of critical HFMD. Over the years, EV-A71 and coxsackievirus A group 16 (CV-A16) were reported as the most common cause of HFMD worldwide, as well as other intestinal viruses including coxsackievirus A group 6 (CV-A6), coxsackievirus A group 10 (CV-A10), echovirus 3, and echovirus 6 [13,14]. Recent epidemiological studies have shown that EV-A71 and CV-A16 have been partially replaced by CV-A6 and CV-A10 as the main viruses related to HFMD; since 2010, there have been several outbreaks of HFMD caused by such viruses in Asia, Americas, and Europe [15,16]. Furthermore, it has been reported that 3.6%–18.2% of severe and critical HFMD were associated with CV-A6 infection during the recent epidemic [17].

Most children with severe HFMD with neurological damage had a good prognosis and high cure rates. However, most critical cases with cardiopulmonary failure had poor prognosis with high morbidity and disability; hence, timely identification of critical cases, especially those with ANS dysfunction, is the key to improve the therapeutic effect of HFMD and reduce mortality [3,4]. Therefore, we screened out the indicators with significant differences among the three groups and then conducted further data comparative analysis between severe group and critical group. Significant indicators between the two groups were EV-A71 positive, heart rate, respiration, cold sweat, moist rales, disturbance in consciousness, and PLT and GLU levels, which were consistent with those reported in previous literature. There was no significant difference in MAP, and the possible reason was the clinical application of vasoactive drugs, which concealed the actual blood pressure of the children with critical HFMD.

In this study, significant differences were found in each scoring system among the three groups. Further logistic regression analysis showed a positive correlation between age-adapted SOFA, qSOFA, PELOD-2, P-MODS, H-SOFA, and critical HFMD, suggesting that the higher the score, the more severe the disease. PCIS was negatively correlated with critical HFMD, suggesting that the lower the score, the more severe the disease. Further ROC curve analysis showed that the AUC value of age-adapted SOFA was 0.938, which was higher than that of age-adapted qSOFA (0.823), PELOD-2 (0.848), P-MODS (0.910), and PCIS (0.865), indicating that age-adapted SOFA had good predictive efficacy for critical HFMD. However, the evaluation of the coagulation system using SOFA was mainly due to the decline in PLT. This study found that the increasing level of PLT was related to severe and critical HFMD. Theoretically, the re-set H-SOFA in our study was considered more suitable for HFMD evaluation. The actual research data also proved that the AUC value of H-SOFA was the highest (0.956), and compared with age-adapted SOFA, it greatly improves the specificity and the prediction efficiency on the premise of no significant loss of sensitivity. Besides, age-adapted SOFA and P-MODS also had good sensitivity and specificity. Age-adapted qSOFA had the highest specificity, but had the lowest sensitivity, resulting in lower predictive efficacy.

In the past decades, HFMD has become more prominent in the Asia-Pacific region [18]. From May 2008 to June 2014, 10,717,283 cases and 3,046 deaths were reported in China, with a fatality rate of 0.03%; among the survivors, the incidence increased from 376/100,000 in 2008 to 1396/100,000 in 2013 [3]. With hundreds of deaths each year, HFMD has become a serious global public health problem. The prevention and treatment of HFMD is still a long way off; the provision of health education helps prevent the transmission, and the promotion of EV-A71 vaccines is of great significance to reduce the severe rate and mortality of HFMD. Early detection, diagnosis, and clinical intervention are of vital importance for children with critical HFMD. The application of PELOD-2, PCIS, and P-MODS in clinical practice, as well as the use of age-adapted SOFA, qSOFA, and H-SOFA will contribute to the recognition of critical cases and enhance the therapeutic effect. However, all studies evaluating the scoring systems are based on population level. Therefore, it is necessary to take into account the huge differences between various individuals. In clinical practice, multiple scoring systems should be applied dynamically for comprehensive evaluation combined with symptoms, signs, and the therapeutic effects. In addition, Shang et al. [19] reported that the plasma expression levels of mononuclear chemokines-1, interleukin-4, interleukin-12, and interleukin-18 were significantly higher in patients with critical HFMD than the mild cases. However, the detection of these indicators has not been carried out in most grassroots medical institutions, which limits the clinical application.

## Conclusions

Increased PLT is associated with the severity of HFMD. The redesigned H-SOFA has a good predictive and evaluation effect on critical HFMD, but its actual application in clinical practice still needs to be further verified by large-scale and multi-center clinical trials.

## Abbreviations

ALB: albumin
ALP: alkaline phosphatase
ALT: aminotransferase
ANS: autonomic nervous system
APTT: activated partial thromboplastin time
AST: aspartate transaminase
AUC: area under the ROC curve
B: Independent variable coefficient
BASO: basophils count
bpm: beats per minute
Ca: serum calcium
CI: confidence interval
CK: creatine kinase
CKMB: MB isoenzyme of creatine kinase
Cl: serum chloride
CNS: central nervous system
CRP: C-reactive protein
CV-A6: coxsackie virus A group 6
CV-A10: coxsackie virus A group 10
CV-A16: coxsackie virus A group 16
DB: direct bilirubin
df: degrees of freedom
EO: eosinophils count
EV-A71: enterovirus A group 71
Fib: fibrinogen
GGT: glutamyl transpeptidase
GLU: blood glucose
HB: hemoglobin
HCT: hematocrit
HFMD: hand foot and mouth disease
H-SOFA: Age-adapted SOFA for Hand Foot and Mouth Disease
IB: indirect bilirubin
IgA: immunoglobulin A
IgG: immunoglobulin G
IgM: immunoglobulin M
K: serum potassium
LDH: lactate dehydrogenase
LYMP: lymphocytes count
MAP: mean arterial pressure
MONO: monocytes count
Na: sodium
NEUT: neutrophils count
NHCC: National Health Commission of China
OR: odds ratio
PB: prealbumin
PCIS: Pediatric Critical Illness Score
PCT: procalcitonin
PELOD-2: Pediatric Logistic Organ Dysfunction Score-2
PLT: platelet
P-MODS: Pediatric Multiple Organ Drgan Dysfunction Score
PT: prothrombin time
qSOFA: quick Sequential Organ Failure Assessment Score
ROC: receiver operating characteristic
SCr: serum creatinine
SE: standard error
SOFA: Sequential Organ Failure Assessment Score
TB: total bilirubin
UA: uric acid
WBC: white blood cell count
WHO: world health organization

## Acknowledgements

The authors would like to thank the staff of the Medical Research Center of Taizhou Hospital for their helpful advice and guidance.

## Funding

This study was supported by the Natural Science Foundation of Zhejiang Province (no. LY16H030001), the Zhejiang Provincial Medical and Health Technology Planning Project (no. 2018KY914), and the Taizhou science and technology planning project (no. 2015A33259 and no. 1801KY22). The funders had no role in the study design, data collection and analysis, decision to publish, or preparation of the manuscript.

## Availability of data and materials

The data analysed during this study are included in this paper. Some of the datasets are available from the corresponding author upon reasonable request.

## Author’s contributions

Z. Yu wrote the manuscript; Z. Yu, A. Li and T. Huang completed the statistical analysis of the data; Z. He, and H. Chen completed the data collection of patients with HFMD; J. Zhu reviewed and edited the manuscript drafts.

## Ethics approval

This retrospective study was reviewed and approved by the Institutional Review Board of the Taizhou Enze Medical Center (Group) Enze Hospital.

## Conflicts of interests

The authors declare no conflicts of interest.

